# Role of slow temporal dynamics in reliable activity of stochastically driven neurons

**DOI:** 10.1101/2020.12.11.422204

**Authors:** Subhadra Mokashe, Suhita Nadkarni

**Affiliations:** Computational Neurobiology Laboratory, Division Of Biology, Indian Institute of Science Education and Research, Pune, India; Neuroscience Graduate Program, Brandeis University, MA, USA

**Keywords:** stochasticity, regular rhythmic activity, CPG, Gillespie algorithm, extrinsic noise, intrinsic noise, reliability

## Abstract

Neuronal networks maintain robust patterns of activity despite a backdrop of noise from various sources. Mutually inhibiting neurons is a standard network motif implicated in rhythm generation. In an elementary network motif of two neurons capable of swapping from an active state to a quiescent state, we ask how different sources of stochasticity alter firing patterns. In this system, the alternating activity occurs via combined action of a calcium-dependent potassium current, sAHP (slow afterhyperpolarization), and a fast GABAergic synapse. We show that simulating extrinsic noise arising from background activity extends the dynamical range of neuronal firing. Extrinsic noise also has the effect of increasing the switching frequency via a faster build-up of sAHP current. We show that switching frequency as a function of input current has a non-monotonic behavior. Interestingly the noise tolerance of this system varies with the input current. It shows maximum robustness to noise at an input current that corresponds to the minimum switching frequency between the neurons. The slow decay time scale of sAHP conductance allows neurons to act as a low-pass filter, attenuate noise, and integrate over ion channel fluctuations. Additionally, we show that the slow inactivation time of the sAHP channel allows the neuron to act as an action potential counter. We propose that this intrinsic property of the current allows the network to maintain rhythmic activity critical for various functions, despite the noise, and operate as a temporal integrator.

---

Several key brain functions critically depend on the reliable activity of neuronal networks. One of the enduring questions in Neurosciences has been to understand how neurons generate robust activity patterns despite an inherently noisy framework. Here we ask how noise arising from intrinsic sources such as thermal fluctuations ion channels and extrinsic sources such as a variable input affects activity in an illustrative network capable of generating rhythms. The network consists of two neurons connected by an inhibitory fast GABAergic synapse that causes neurons to switch off as synaptic current builds up (See Figure 1). Mutually inhibitory networks of neurons -are a recurring motif across brain areas, for example; in the hippocampus (Pelkey et al., 2017), central pattern generators associated with locomotion and digestion (Otto Friesen, 1994), insect olfactory systems (Daun et al., 2009), REM sleep cycle (Lu et al., 2006), and working memory (Myre and Woodward, 1993). The on-off switching activity of neurons allows them to be associated with multiple networks (Hooper and Moulins, 1989). It dictates sequential order of activity required, for example, locomotion (Cangiano and Grillner, 2004) and spatial navigation Dragoi and Buzsáki (2006).

**Figure 1.**
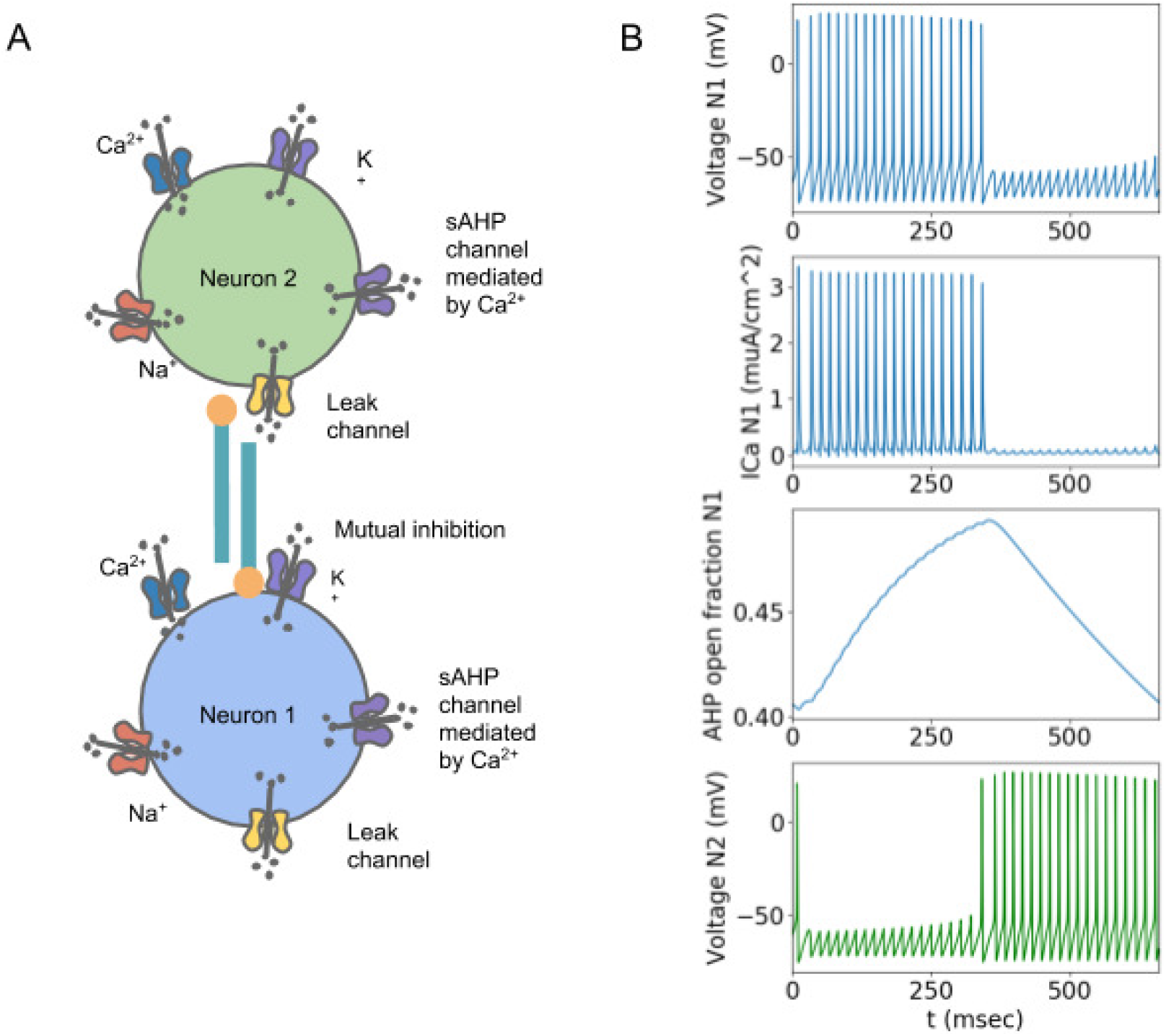
Network and switching mechanism: A. The network of mutually inhibiting neurons along with the ion channels which orchestrate the firing and switching activity of the neurons. B. Top to bottom: Neuron fires action potentials due to depolarization driven by the external current. Calcium channels open a result of depolarization of the membrane leading to calcium ion influx. The build-up of sAHP current over multiple action potentials leads to termination of burst in neuron 1. Escape from inhibition and burst of the neuron 2.

In the system described in Figure 1, switching occurs due to an action potential (AP) triggered build-up of a calcium-dependent potassium current, specifically called slow after-hyperpolarization (sAHP). The potassium current causes the neuronal membrane to be hyperpolarized. This hyperpolarization is long-lasting due to the intrinsic slow closing time of the sAHP channel. As the potassium current builds up, it makes it incrementally harder for the neuron to fire an AP in response to a stimulus (conductance of sAHP grows with every calcium spike that closely follow APs). Eventually, the neuron unable to generate an action potential, deactivates the inhibitory synaptic current (GABAergic) to the connected neuron. This release from inhibition enables the other neuron to fire action potentials, creating the 121212 activity pattern. A detailed chronology of events is as follows: Strong DC input is applied to both the neurons (with identical intrinsic properties but slightly different initial conditions). The neuron with advantageous initial conditions takes over and generates action potentials. The resulting synaptic current inhibits activity in the other neuron via a GABA synapse. In the active neuron, in the meantime, APs cause the opening of voltage-gated calcium channels (VGCCs). VGCCs follow the open-close action of APs closely. The incoming calcium flux through VGCCS activates the sAHP channels (Figure 1 B top three panels). sAHP channels respond rapidly to calcium. However, as the name suggests, sAHP channels inactivate slowly; therefore, with each calcium pulse, the number of active sAHP channels increases. The slow closing time of the channels allows them to remain open even after calcium channels are closed and calcium is extruded out, ensures that potassium builds over multiple action potentials. Beyond a hyperpolarization threshold induced by the potassium current, the neuron is disabled. Thus the action of sAHP terminates the AP activity after a characteristic time interval governed by the potassium current build-up (Manira et al., 1994)(Figure 1 B top third panel). The termination of the burst of APs puts an end to the active inhibitory synapse. The other neuron gets activated now and completes the rhythmic pattern(Figure 1 B bottom panel).

The overall time over which a single neuron in this network remains active is a function of inactivation time constant of the sAHP current and synaptic current, stimulus strength, calcium ion flux, intrinsic noise due to fluctuations of ions channels, and extrinsic noise arising out of modulation of the stimulus (Figure 1 A) (Figure 1 B third top panel). These contribute to the potassium current differentially and dictate the burst interval.

Effects of various sources of noise on system behavior and, generally on brain function have been extensively investigated (Goldwyn and Shea-Brown, 2011). Noise can be both disruptive and enhance function (Stacey and Durand, 2001). The addition of noise can increase signal detection and transduction via stochastic resonance in Hippocampal CA1 neurons. (McDonnell and Abbott, 2009; Schmid et al., 2001; Stacey and Durand, 2001). Coherence resonance is another interesting phenomenon that arises due to noise but increases the regularity of activity (Andreev et al., 2018). Apart from these external sources of noise (extrinsic noise), intrinsic sources of variability such channel fluctuations can also modify function (Schmid et al., 2001). Noisy opening and closing of voltage-gated calcium channels can allow intracellular calcium influx and trigger downstream calcium-mediated signals, make the neuron more excitable, allow transmission of subthreshold signals (White et al., 2000) and cause a post-inhibitory rebound effect (Tegnér et al., 1997).

Here we systematically investigate the consequences of significant sources of intrinsic and extrinsic noise on the reliability of switching in an essential functional network capable of rhythmic behavior: two neurons connected by an inhibitory synapse (Figure 1). The effect of extrinsic noise is studied by varying the amplitude of the current noise. It has been shown that voltage-gated calcium channels (VGCCs) fluctuations are one of the main contributors to the stochasticity at the synapse (Modchang et al., 2010). We study the influence of intrinsic noise arising from channel fluctuations of the VGCCs. We ask how calcium channel noise modulates sAHP conductance and, in turn, changes the switching rate.

## 1 METHODS

We used a ionic conductance-based model of neurons that are connected to each other via an inhibitory synapse. The potassium current, sAHP (slow afterhyperpolarization) which is mediated by calcium ions orchestrates switching in the network (equations (1), (2)). Extrinsic noise is interpreted as the noise arising independently of the state of the neuron, such as the background noise, and is implemented here as an additive term *ξ* to the differential equation of the voltage (equations (1), (2)). In contrast, intrinsic noise depends on the state of the neuron. It is implemented in the model as the stochasticity associated with a small number of ion channels and stochastic channel opening. To model realistic intrinsic noise, we simulate a Markovian description of the calcium channels using the Gillespie algorithm.

### 1.1 Network model

The neurons have voltage-gated calcium channels, and sAHP channels along with voltage-gated sodium and potassium channels leak current and extrinsic noise.

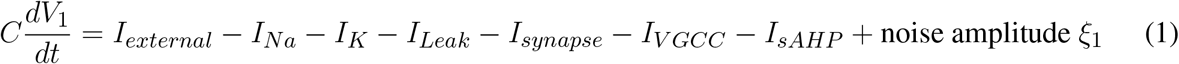

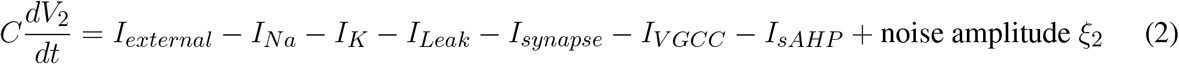

### 1.2 Hodgkin-Huxley Neuron Model

The classical Hodgkin-Huxley neuron model describes how neurons generate action potentials (Hodgkin and Huxley, 1990). It has *Na*^+^, *K*^+^, and leak channels given by,

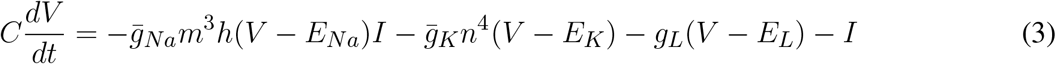

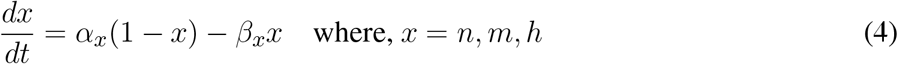

Where,

*V* : membrane potential

*n, m, h*: gating variables which represent the open fraction of channels of sodium (*m, h*) and potassium(*n*). C = 1 *µ*F/*cm*^2^: the capacitance of the cell membrane

*E*_*Na*_ = 50 mV, *E*_*K*_ = −77 mV, and *E*_*L*_ = −54.4 mV: reversal potentials of sodium, potassium and leak channels respectively.

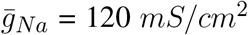 and 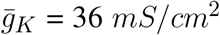: maximal conductances of sodium and potassium currents respectively.

*g*_*L*_ = 0.3 *mS/cm*^2^: leak conductance

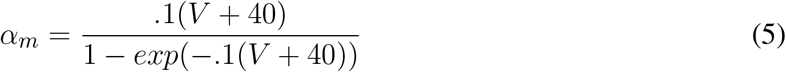

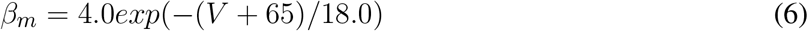

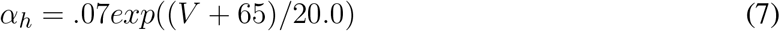

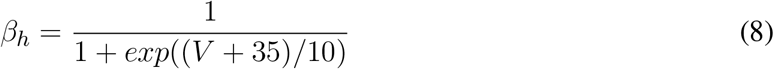

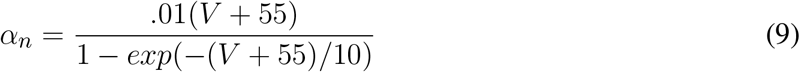

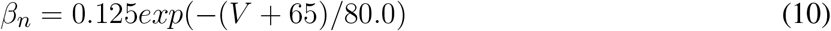

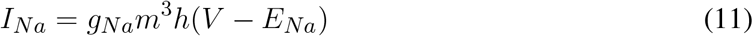

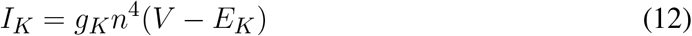

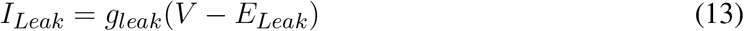

### 1.3 sAHP channels

The model for calcium-mediated potassium current, sAHP is based on the sAHP channels of CA1 pyramidal neurons (Sah and Clements, 1999) (Stanley et al., 2011).

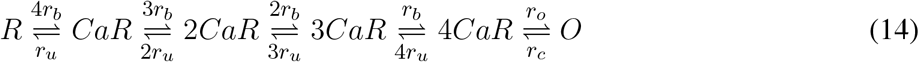

Where *r*_*b*_ = 4 *µ* M/sec, *r*_*u*_ = 0.5/sec, *r*_*o*_ = 600/sec, and *r*_*c*_ = 400/sec. Here R, CA1R, CA2R, CA3R, CA4R, and O are the states of the channel. R is the closed state, and O is the open state. The total conductance of the channel is dependent on the fraction of open channels. The peak open probability of the channel is 0.4, and its mean open time is 2.5 msec. When [*Ca*^2+^]_*i*_ falls rapidly, the decay of sAHP is limited by the channel closing and *Ca*^+2^ dissociation rates to give a time constant of 1.5 sec (Sah and Clements, 1999).

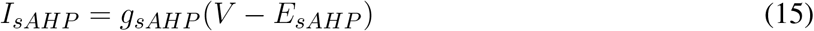

Where *g*_*sAHP*_ = 0.4 *µ*S/*cm*^2^ and *E*_*sAHP*_ = *E*_*K*_ = −77 mV

### 1.4 Synapse

Inhibitory synapses are modelled using a tan hyperbolic function.

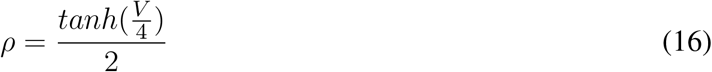

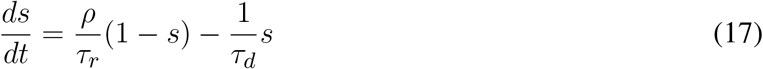

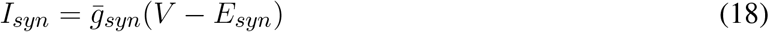

Where 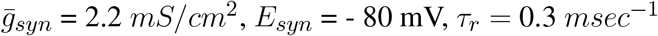 and *τ*_*r*_ = 8.9 *msec*^−1^ .

### 1.5 Voltage gated calcium channels

We model the L-type *Cav*_1.3_ calcium channels which open at low voltages given by (Stanley et al., 2011).

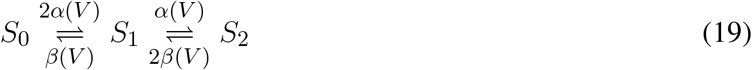

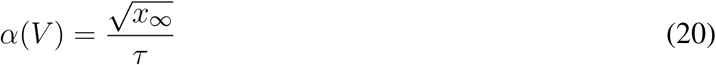

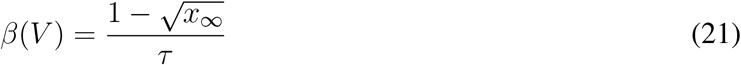

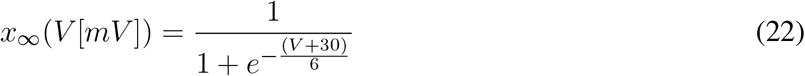

Here *α, β* are voltage dependent probabilities of transitions of states *S*_*i*_ . The conductance is dependent on the fraction of open state.

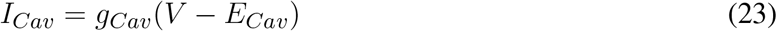

Where *g*_*cav*_ = 0.15 mS/ *cm*^2^ and *E*_*cav*_ = *E*_*Ca*_ = 25mV

### 1.6 Modelling calcium dynamics

The intracellular calcium concentration dynamics is modeled as a leaky integrator, (Stanley et al., 2011) (Wang, 1998).

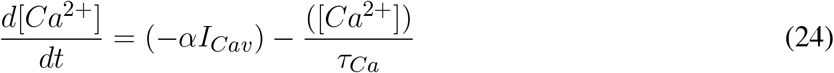

Where *α* = 2.10^−4^[*mM* (*msecµA*)^−1^*cm*^2^] and *τ*_*Ca*_ = 14 ms. *α* depends on the area to volume ratio of the neuron, intracellular buffering of calcium, and stochasticity factor, and converts calcium current into the units of calcium concentration per unit time. The resting calcium concentration is 100 nM and goes up to 2.5 *µM* per spike.

### 1.7 Modelling channel noise

For large channel numbers, the fluctuations in the conductance of channels are small and can be modeled using deterministic dynamics. However, a small number of ion channels typically dictate the neuronal dynamics. Under these circumstances, the stochasticity of ion channel fluctuations becomes relevant. Channel noise has been extensively studied, and various methods to model channel noise have been explored (reviewed in (Goldwyn and Shea-Brown, 2011). Stochastic dynamics simulated using the Gillespie algorithm is a fast and accurate algorithm to simulate channel noise (Gillespie, 1976b).

Given that the fluctuations arising out of VGCCs are significant contributors to noise in the calcium signal that ultimately governs switching dynamics, we selectively target the investigation of noise arising from calcium channel fluctuations. Towards this control experiment, we implement Markovian description only VGCCs using the Gillespie algorithm (Gillespie, 1976a), whereas the other components of the model are modeled deterministically. To accurately capture all transitions, We developed an algorithm to implement the Gillespie algorithm (for Markovian progression) and Euler method (for deterministic progression) in tandem for a system of equations with multiple timescales that span several orders of magnitude ((Stanley et al., 2011) (slowAHP *τ* = 1.5 s, fast voltage, and calcium dynamics *τ* =14 ms). We call this Gillespie-Euler Hybrid Algorithm, ‘Tandem Progression Gillespie (TPG), used to simulate realistic time scales and amplitudes of channel noise.

In our implementation of the Gillespie Algorithm, we updated the entire system at 1) the fixed time step dictated by the deterministic part of the model and 2) the waiting times obtained from modeling the calcium channel dynamics as a Poisson process. This is distinct from Chow and White (1996), where the voltage is updated only at times dictated by channel transitions. This modification was crucial as the build-up of sAHP due to calcium fluctuations would be missed between the channel waiting times otherwise. This would be especially true when the waiting times are longer. Thus by updating the whole system together at a time that arises either out of the calcium channel fluctuations or voltage equations, the dynamics of this multi-timescale system evolved more accurately via TPG.

In the algorithm by Goldwyn et al. (Goldwyn and Shea-Brown, 2011) (Model DB accession number 138950), voltages are updated at a fixed time step. This algorithm is correct under the assumption that 1) The rates for transitions do not change between two time-steps and 2) There are no slow timescales involved which could keep track of all the fluctuations, as the fluctuations between the fixed time steps may not be seen by the voltage and other currents in the neuron.

#### Algorithm 1. Goldwyn and Shea-Brown Gillespie Modification

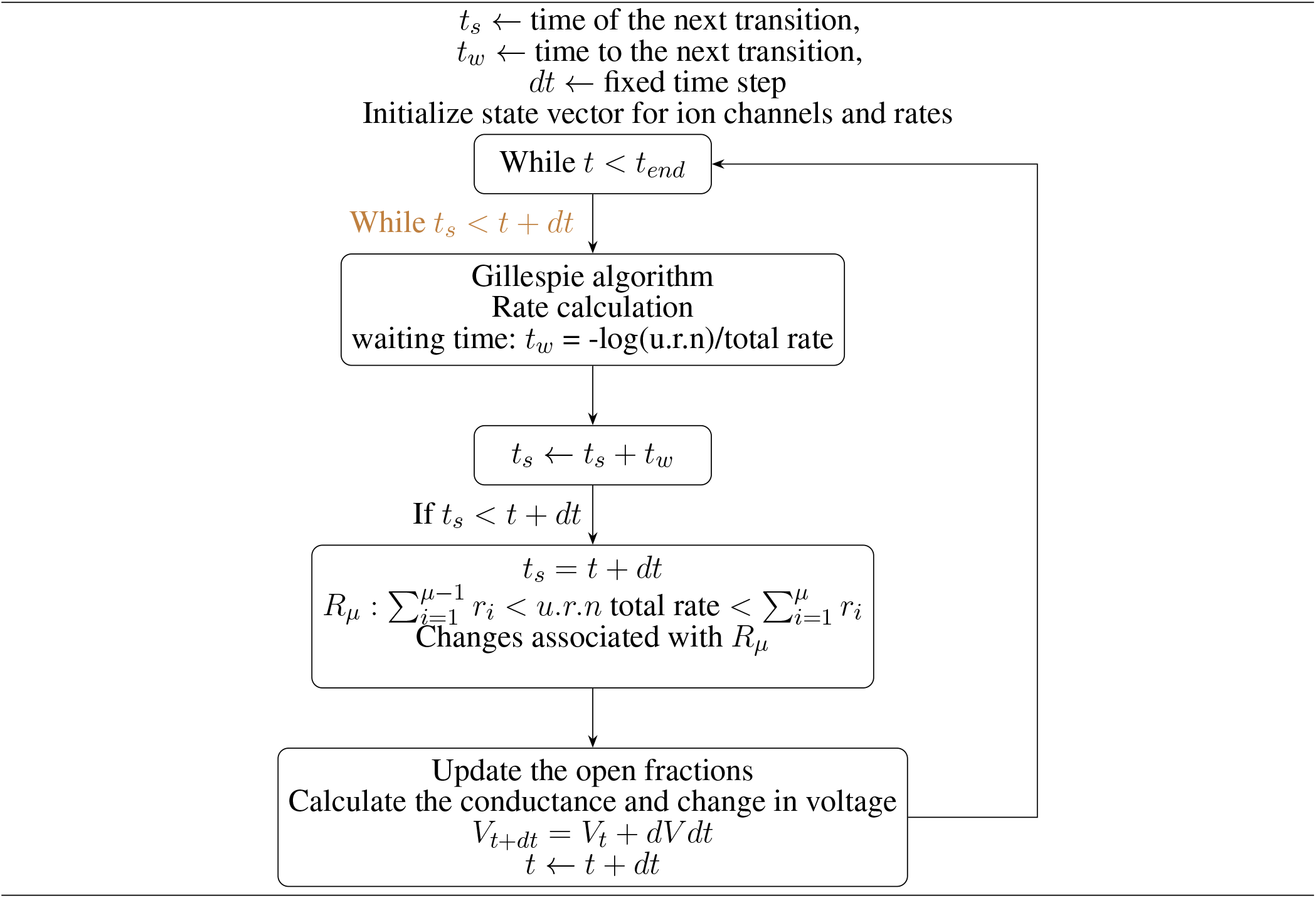

Another implementation of the Gillespie algorithm for conductance-based neurons suggested by Chow et al. (Chow and White, 1996) integrates the deterministic system till the waiting time given by the Gillespie algorithm and updates the stochastic system only after Gillespie waiting times. This algorithm assumes that the rate constants do not change during the time step, and the rate and waiting time calculations take into account the dynamics and time-scales from all the ion channels present in the neuron. Yet another approach used to model channel fluctuations: the system size expansion approach used by (Fox and Lu, 1994), which involves solving the drift-diffusion equation to accurately model the stochastic dynamics simulated using the Gillespie algorithm since the Gillespie algorithm is computationally expensive. This approach was not appropriate as the system is not large enough and also would compromise accuracy.

To isolate the influence of noise due to VGCC fluctuations, we use the Markovian kinetic scheme to simulate channel dynamics, whereas the rest of the system is allowed to evolve deterministically. Since we wanted to study the effect of the channel noise arising from one type of ion channel, these modifications were essential. Integrating at fixed time steps as in (Goldwyn and Shea-Brown, 2011) will lead to missing channel fluctuations that take place between two time-steps. The fast activation and slow decay timescales associated with the sAHP current will cause the fluctuations caused by a noisy current input to have a cumulative effect on the sAHP current and significantly modify switching times between neurons. In order to not miss these fluctuations, we update the whole system deterministic and in ‘tandem’, the stochastic system at the Gillespie algorith determined time-steps. While integrating merely at long waiting times associated with small channel numbers, the dynamics of the other components of the model neuron may not be captured correctly and could lead to errors. To model the neuronal dynamics correctly, especially when the waiting times are longer than a fixed time step (0.01 msec used in simulations), we integrate the system at the fixed time step and also update the stochastic channel states at every integration step to take into account the changed voltage and current values. Thus by updating the whole system together, we believe that we are modeling the stochastic channel dynamics as well as the neuronal and network dynamics accurately. In summary, in Tandem Progression Gillespie, every component of the model is updated at the same time and is described in Algorithm 3.

#### Algorithm 2. Chow and White Gillespie Modification

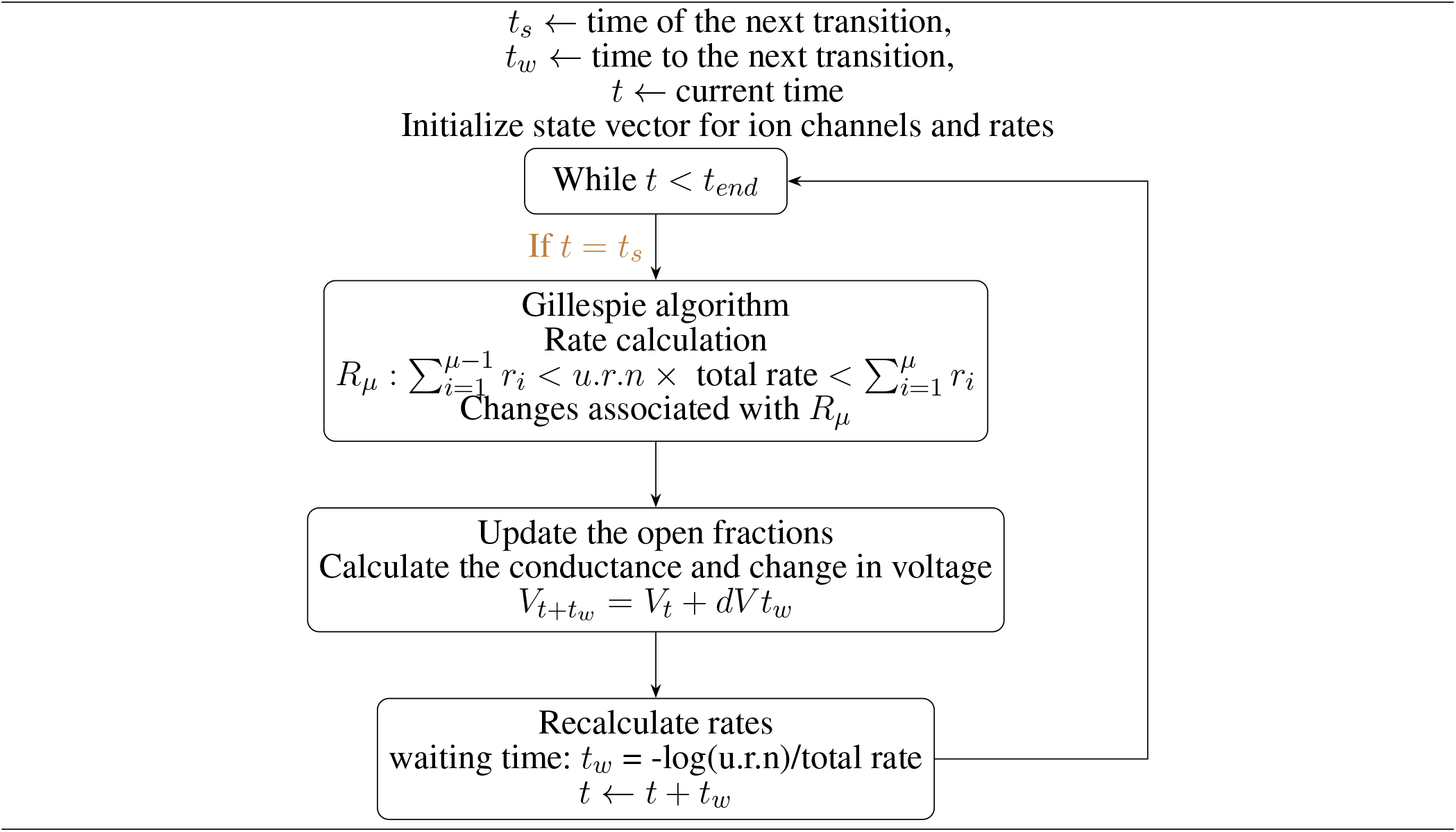

For higher channel number, Chow and White and TPG algorithms show similar trends in switching, see in Figure 2 as most transitions occur at the Gillespie waiting time.

**Figure 2.**
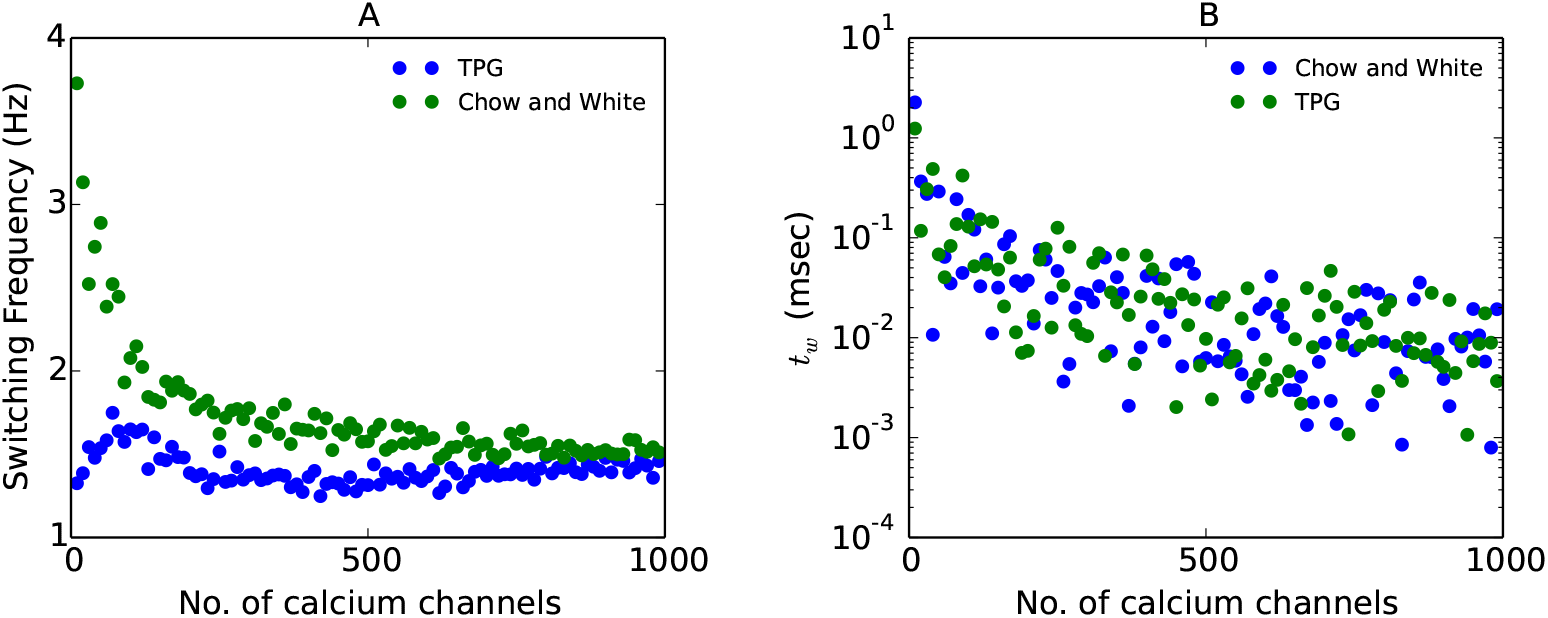
Comparision of switching dynamics for TPG and algorithm used in (Chow and White, 1996) A. The mean switching frequency described by TPG, and, Gilespie implementation of Chow and White . B. The CV of interburst intervals for different numbers of calcium channel for the two algorithms.

### 1.8 Calcium channel opening failures

To test how sAHP integrates over irregular and unreliable calcium signal, we induce calcium channel opening failures with a given probability. Calcium failures are modeled either as individual channel failures or as ensemble level or pulse failures. Ensemble level failures are calcium pulse failure. Each calcium pulse can be invisible to the neuron and thus sAHP current with a certain probability (failure rate). The calcium current comes up again after the failed calcium pulse, and the failure is limited to the duration of calcium pulse and is carried out by multiplying a voltage-dependent block on the calcium current. In this case, all channels fail to open during the block. In the case of an individual channel failure, each channel opening transitions fail with a certain failure probability (failure rate), and thus only one channel fails to open.

#### Algorithm 3. Tandem Progression Gillespie

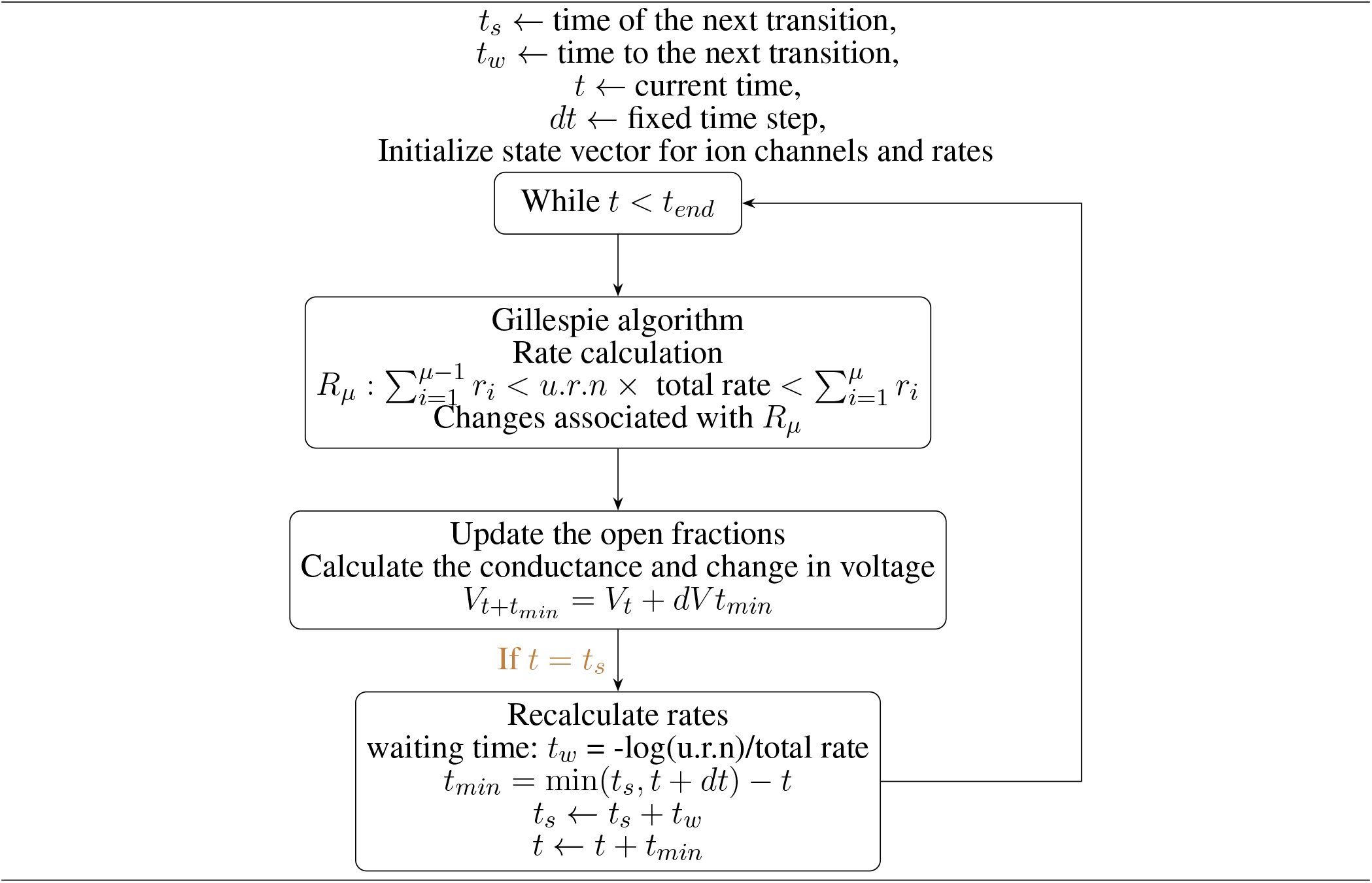

### 1.9 insilico

insilico is a C++ based computational tool specifically designed developed to simulate neurons. The deterministic model is implemented using insilico-0.25 http://www.iiserpune.ac.in/~collins/insilico/.

### 1.10 Analysis

To study the effect of various parameter on the switching dynamics, we look at the burst length and the switching frequency of the neurons which is the primary functional read-out of the network.

#### Inter-spike interval and Firing frequency

The time difference between peaks of two consecutive action potentials is the inter-spike interval, and the inverse of the inter-spike gives us the firing frequency of the neuron.

#### Inter-burst interval and switching frequency

We define switching frequency as the frequency with which the neurons alternate in their activity. Burst is defined as a set of action potentials the neuron fires before the other neuron is released from inhibition and starts firing When the burst terminates because of sAHP current, and other neuron takes over and inhibits the first neuron, the interval between the last action potential from the last burst to the first action potential of the next burst of the neuron is called an interburst interval(IBI). The inverse of IBI is called switching frequency or burst frequency.

The action potential is detected if the voltage goes higher than 15 mV and if such a detection happens after a minimum of time difference of 5 msec after the last detection. A burst is detected when interspike intervals greater than twice the last inter-burst interval. We calculate the switching frequency by finding the inverse of the mean of a fixed number of burst lengths. As a measure of regularity of bursts, coefficient of variation is calculated where *T* is the inter-burst interval, is given by,

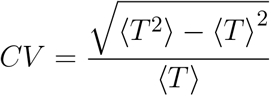

## 2 RESULTS

### 2.1 Modulation of switching frequency by driving current

The total time taken for the activity to switch from one neuron to the other, called the ‘Inter-Burst-Interval’, depends on external current stimulus *I*_*ext*_, its influence on synaptic conductance, *G*_*syn*_, the time constant of the synaptic current *τ*_*syn*_, the conductance of the sAHP current *g*_*sahp*_, and the AHP-calcium-binding rates. For physiologically realistic synaptic coupling strengths and sAHP conductance (*g*_*sahp*_ = 0.4*µS/cm*^2^, synaptic conductance = 2.2*µS/cm*^2^) (Sah and Clements (1999)), the switching of activity between neurons is modulated over a couple of Hz (0.8 Hz-3 Hz) and observed for an external current between 13 *µA/cm*^2^ to 18.75 *µA/cm*^2^. Switching ceases beyond this range of external current. The dependence of switching frequency on the external current can be summarized as follows; An increase in driving current makes both the neurons more excitable and leads to an increase in spiking frequency (Figure 3 A). This increase leads to an increased rate of calcium spikes and a faster build-up of sAHP current, which can shorten the duration over which the neuron is active (burst duration). On the other hand, the increase in depolarizing input drive also increases the hyperpolarization needed to terminate the burst via the sAHP current (expressed in terms of threshold sAHP open fraction) (Figure 3 B). This has the effect of increasing the duration of the burst, as it takes longer to reach the threshold sAHP current. The increase in the threshold of the sAHP current needed to terminate the burst is shown in Figure 3 B. These distinct opposite effects on the rate of sAHP built-up in response increase in driving current are shown in Figure 3 C. Enhanced depolarization dictates the initial decrease in the rate of AHP_threshold_ until *I*_*ext*_=16 *µA/cm*^2^. It is followed by an in the rate of the sAHP build-up due to a faster rate of incoming calcium spikes. The initial the spike frequency increase causes a decrease in the switching frequency (Figure 3 D). It can be explained by the time taken for sAHP to achieve higher conductance levels thus extending the switching time. Between the input current 15 *µA/cm*^2^ and 16 *µA/cm*^2^, the frequency of switching remains fixed at 0.8 Hz. However, beyond 16 *µA/cm*^2^, the switching frequency increases as it follows the sAHP build-up rate. Increasing external driving current to both the neurons described by *I*_*ext*_ in equation (1) (while keeping other parameters unchanged), thus, has a non-monotonic effect on the switching frequency (Figure 3 D).

**Figure 3.**
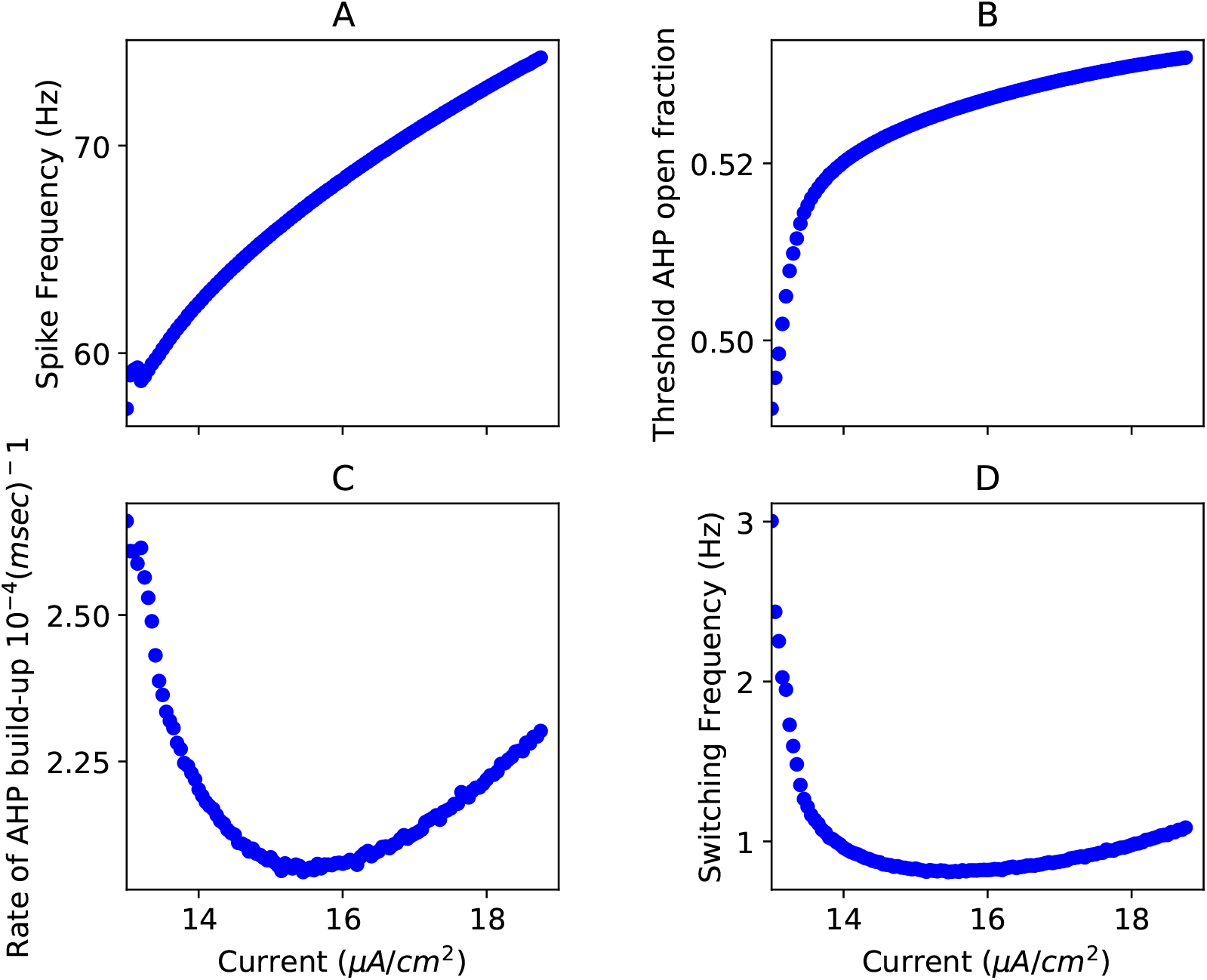
Frequency modulation by current: Competition between an increase in excitability and sAHP threshold current leads to non-monotonic changes in switching frequency: A. The spike frequency increases with increasing current.B. The peak sAHP open fraction reached increases monotonically on increasing the current. C. The rate of build-up sAHP current shows a minimum at an intermediate current value. D. The switching frequency shows a non-monotonic dependence on the external current due to two opposing effects that increasing current has on the neuron’s excitability.

### 2.2 Modulation of switching frequency by extrinsic noise

To investigate the effect of extrinsic noise on the switching times between the two neurons, we simulate additive current noise (*ξ*, equation (1)). Recall that regular switching is orchestrated by calcium current and intrinsic opening and closing timescale of the sAHP current. At intermediate current values, a minimum in the coefficient of variation (CV) of switching response (Figure 4 A, blue) is observed that corresponds to a minimum in sAHP rate (Figure 4 A, green). This indicates that the network is most insensitive to external noise for a finite range of input current. At these values of intermediate current, the slow AHP current integrates noisy input current best to maintain regular switching. At high input current and the consequent stronger depolarization, a higher threshold AHP fraction is needed to achieve termination of the burst. This makes the burst duration longer and an improved smoothening of the noisy input by the sAHP current. The filtering of noisy input is seen as a lower CVs and (see Figure 4 A blue) the cumulative effect of the fluctuations is seen as an increased switching frequency.

**Figure 4.**
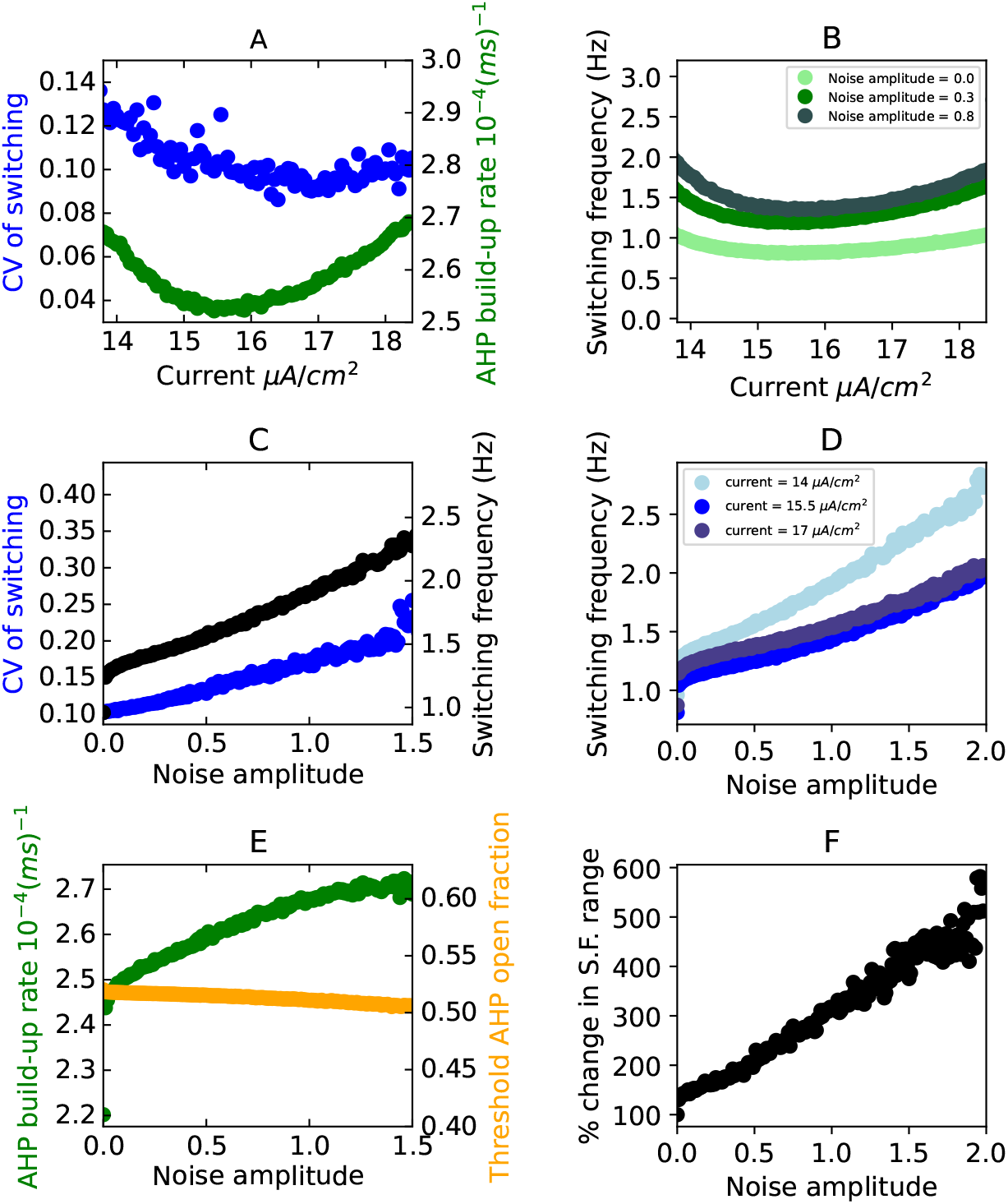
Optimal excitability helps in maintaining reliable switching in the presence of extrinsic noise: A. The minimum in the sAHP build-up (green) rate corresponds to a minimum in the CV of switching (blue) for noise amplitude 0.8. B. The switching frequency increases on increasing the amplitude of the noise. C. The switching frequency (black) and the CV of switching (blue) increase on increasing the noise amplitude for current = 14*µA/cm*^2^. D. Switching frequency versus the noise amplitude for three driving current values. E. The rate of sAHP-build (green) increases as the noise amplitude is increased. In contrast, the threshold sAHP conductance (open fraction in yellow) remains almost constant on increasing the noise amplitude for current = 14*µA/cm*^2^. F. The switching frequency increases by ∼350 percent for current = 14*µA/cm*^2^ with the addition of noise along with some unreliability in switching (CV=0.2 at noise amplitude =1.5).

We show that switching frequency (Figure 4 B, C (black) and D)increases with noise amplitude. The switching frequency for three illustrative current values as a function of noise amplitude is shown in (Figure 4 D).

The role of sAHP current in increasing the switching frequency in response to noisy inputs can be understood in the following way: Current noise causes voltage fluctuations leading to fluctuations in the voltage-gated calcium current. The sAHP current can note each of these fluctuations in calcium concentration due to its fast rise-time. However, as the sAHP current’s decay-time is long, the rise in sAHP due to calcium fluctuations accumulates over time and leads to a quicker build-up of sAHP current (Figure 4 E), green). The faster build-up of sAHP terminates the burst earlier, increasing the switching frequency (Figure 4 C, black) with a minor increase in the CV of switching (Figure 4 C, blue). The stochastic calcium signal is filtered with a slow decay-time of the sAHP current and only the summation over time rather than the individual calcium fluctuations dictate the termination of the burst. The increase in switching frequency mediated by noise goes up with amplitude of input current (Figure 4 D).

In figure 4 F, the percentage change in switching frequency range (calculated as: 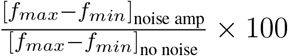) is shown. The addition of noise increases the range of switching frequencies achieved by ∼350 % at noise amplitude 0.2. In summary reliable switching over a wider range of frequencies can be achieved by input noise due to the biophysical properties of the sAHP current.

### 2.3 Modulation by intrinsic noise: effect of calcium pulse failures on switching dynamics

To further examine the sensitivity of sAHP response to the fluctuations in calcium current, we introduce random calcium pulse failures at a varying rates. Increasing the rate of calcium pulse failure decreased the switching frequency (Figure 5 A) due to a slower build-up of sAHP current (increasing number of calcium pulses are missed by sAHP current). Predictably, the increase in the failure rate has the effect of increasing the CV of switching (Figure 5 A). In the presence of a “stochastically faulty” calcium pulse generator, more action potentials need to be fired by the neuron to achieve the threshold sAHP current to terminate the burst. We show the increase in the failure rate (probability) leads to an increase in the number of action potentials per burst (Figure 5 B). Needing more action potentials to achieve the burst termination also makes the burst longer. Thus switching frequency decreases with the increase in the number of action potentials in the burst (Figure 5 C). A linear decrease in switching frequency with increasing the failure rate (Figure 5 A) is seen. Interestingly, similar trends are seen for independent random trials of the calcium pulse failures occurring randomly during the epoch of the burst. The invariance to the temporal position of missed calcium pulses during the burst indicates that the switching behavior is dictated by the number of calcium pulses causing the build of sAHP current and is insensitive to the temporal order of these calcium pulses.

**Figure 5.**
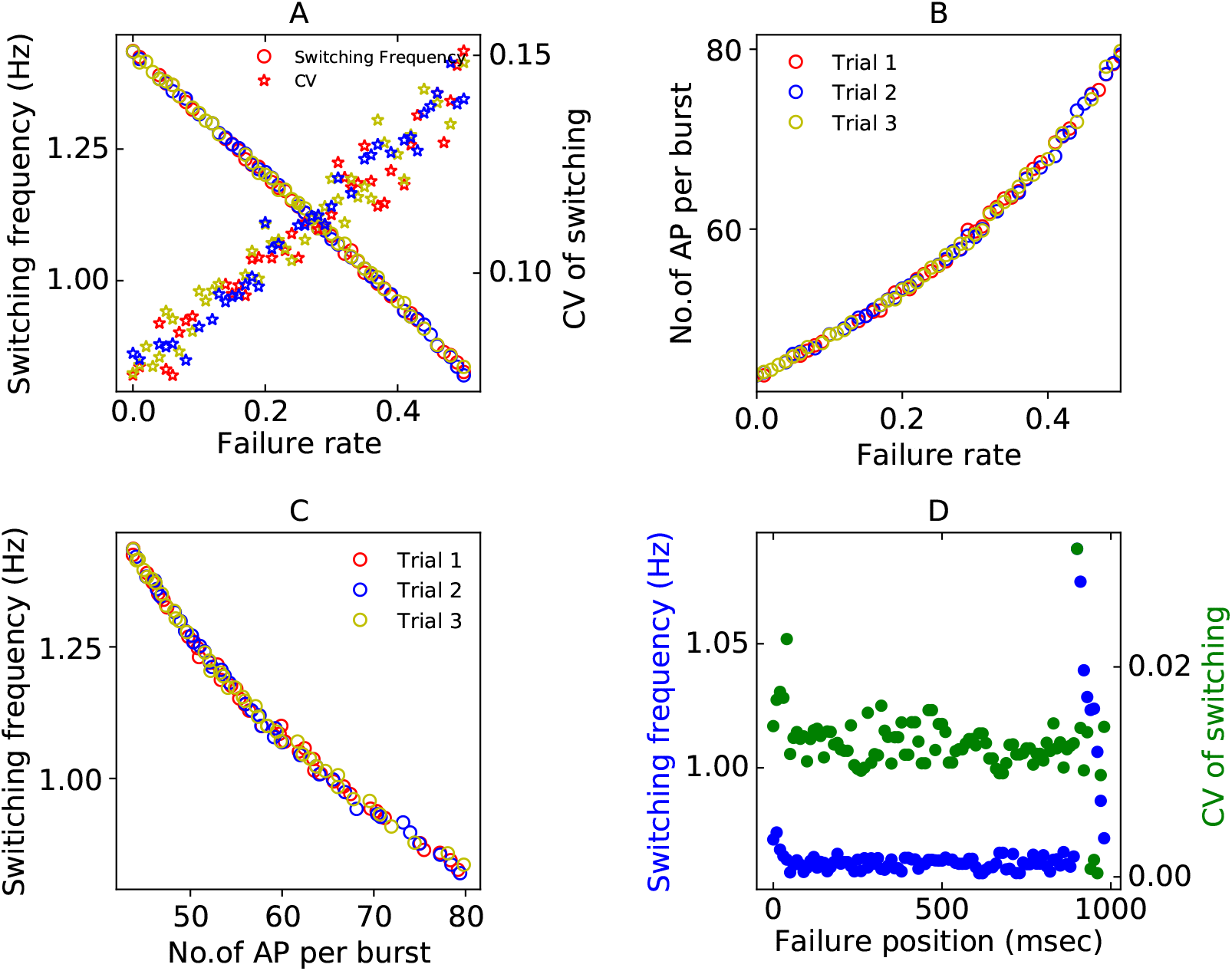
Switching dynamics in the presence of calcium pulse failures. A. The mean switching frequency(circles) and CV of switching(stars) for changing three different trials of calcium pulse failures for a range of failure rates. B. Average number of action potentials in a burst as a function of failure rate. C. The Switching frequency is a function of failure rate and correlates negatively with the number of APs per burst. D. One single calcium pulse failure is induced, and the position of failure is varied during the burst (failure position zero indicates that the failure happened as the initiation of the burst and failure position 1000 indicates that the failure happened after 1000 msec of the burst initiation.) Switching frequency (blue) and CV (green) of switching as a function of failure position.

To systematically test the insensitivity of the sAHP current to the temporal characteristic of the stimulus train, we induced one failure per burst. We changed the position of the calcium pulse failure relative to the initiation of the burst. We see that the position of the failure does not affect the switching frequency for most of the burst and only affects towards the end of the burst(above 900 msec in failure position) (Figure 5 D, blue). The failure at the end of the burst has a drastic effect on the switching frequency. At the end of the burst, the depolarization is weak, and the inhibition from the inhibiting neuron is strong. Thus missing a calcium pulse close to the termination of the burst reduces the depolarizing current to the neuron. In the meantime, the other neuron takes over and inhibits the neuron whose burst is about to end. This results in increased switching frequency when calcium failures appear at the end of the burst (Figure 5 D blue). A missed calcium pulse here also increases irregularity in switching (Figure 5 D, green). However, the CV of the switching remains the same for most missed temporal positions of the calcium pulse. This is illustrated in Figure 5 D, green. In summary, the sAHP current serves as an action potential counter and makes the neuron insensitive to the variations in temporal patterns of the stimulus.

### 2.4 Modulation of switching frequency by calcium channel noise

The slow closing times of the sAHP channels result in the sAHP current maintaining a long memory of these fluctuations. The stochastic opening of calcium channels is the most significant contributor to the variability in synaptic release (Modchang et al., 2010). In order to isolate the effect of calcium channel stochasticity on switching, markovian opening and closing of channels were simulated; however, the rest of the system equations were simulated deterministically. Recall that our two neuron system has intrinsic timescales that vary over a wide range (over two orders of magnitude *τ*_*sAHP*_ = 1.5*sec, τ*_*cal*_ = 14*msec* (Stanley et al., 2011)). The extant algorithms for Markovian simulations; Gilespie algorithm (Gillespie, 1976b) and the (Goldwyn and Shea-Brown, 2011) introduce errors that accumulate with time due to the intrinsic differences in timescales. We, therefore, developed an updating system for variables that we call the Tandem Progression Gillespie or TPG algorithm (See methods for details). According to TPG; 1) transitions in the calcium channel opening get updated according to the Gillespie algorithm and 2) the deterministic changes in the rest of the variables (activation and inactivation gates of the ion channels) get updated according to an Euler integrator. Thus by updating the whole system together at a time that arises either out of the calcium channel fluctuations or voltage equations, the dynamics of this multi-timescale system evolved more accurately.

We see that the switching frequency follows a non-monotonic trend with increasing the number of VGCCs (Figure 6 A). The CV of switching decreases with increasing channel numbers (Figure 6 B) as the fluctuations become smaller in amplitude when the number of calcium channels is increased (Figure 6 C). Beyond about 100 VGCCS, the switching frequency decreases as the number of VGCCs is increased. The waiting times from the Gillespie algorithm become shorter as the number of VGCCs are increased (Figure 6 D). When the number of VGCCs is further increased, beyond 400 VGCCs, the fluctuations become smaller in amplitude, but their frequency increases (Compare Figure B and D). Due to the slow timescales of the sAHP current, these closely spaced fluctuations add up in sAHP current leading to a faster build-up of sAHP current resulting in the increase trend in switching frequencies at large numbers of VDCCs. Thus the increasing the calcium channel numbers causes a) decrease in the amplitude of fluctuations, b) an increase in the frequency of fluctuations, which have opposing effects on the switching frequency. The two opposing effects cause the non-monotonic trend is observed in the switching frequency upon increasing the number of calcium channel numbers. These simulations are carried out for the same maximum conductance of the calcium current.

**Figure 6.**
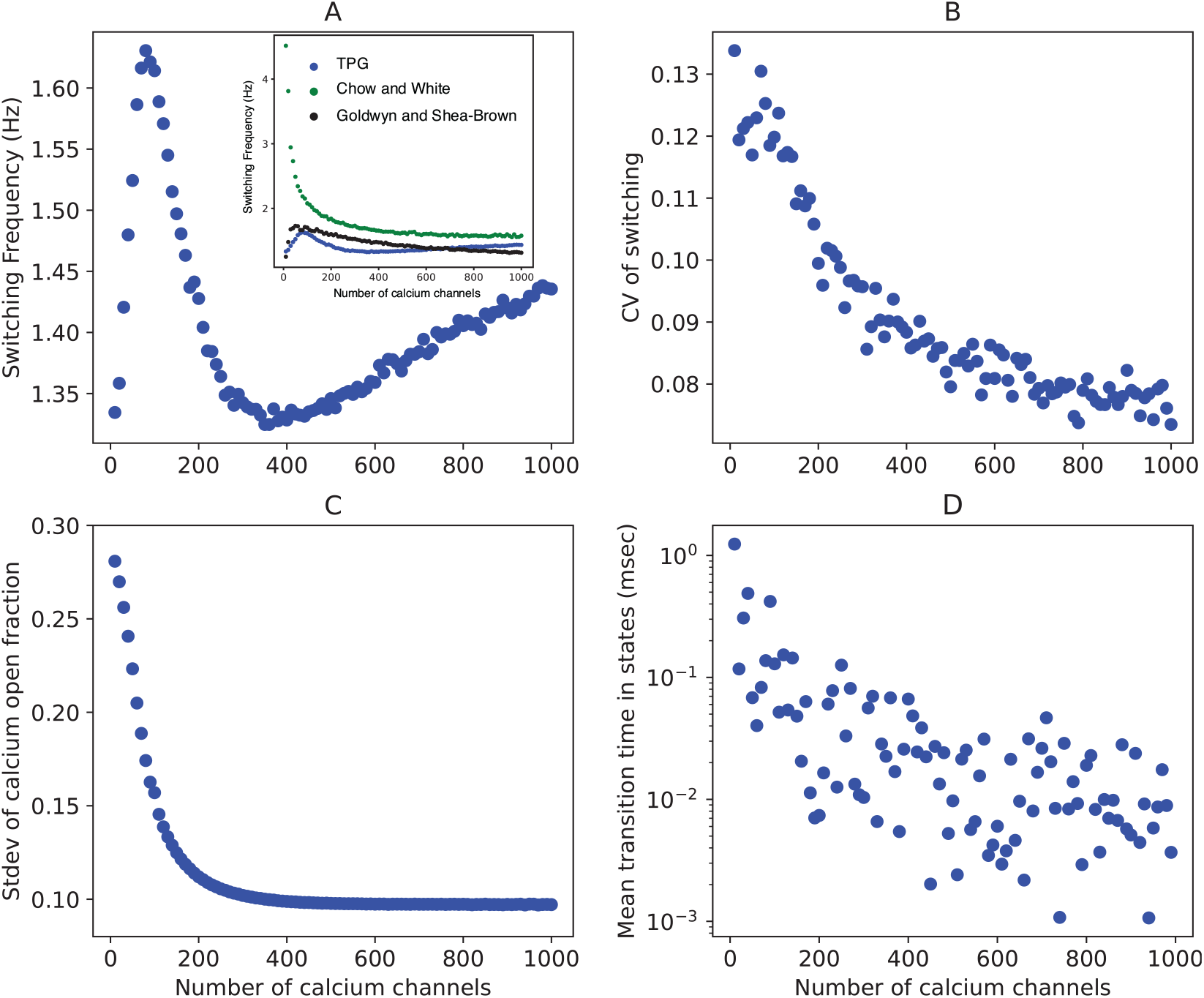
Opposite effects of increasing the number of channels leads to non-monotonic variation in switching frequency. A: The switching frequency shows a non-monotonic trend with increasing the number of calcium channels. B. The switching becomes more regular as the amplitude of fluctuations decreases with the increasing of the number of calcium channels. C. The fluctuations in the open fraction of calcium channels become smaller in amplitude with the increasing channel number. D. The mean transition time in the calcium channel states decreases as channel number is increased, leading to increased frequency of fluctuations in calcium concentration.

## 3 DISCUSSION

It is conjectured that the brain is fundamentally a rhythm generating machine (Buzsáki, 2006). All functions of the brain, from motor patterns, breathing to cognition, emerge from various rhythms. Reciprocal inhibition between neuron pairs is ‘the’ building block that generates stable anti-phasic and multiphasic output patterns. The switching in these reciprocally inhibited networks occurs due to either intrinsic biophysical properties of ionic currents that assign a distinct periodicity to the rhythm or an external drive. There is a distinct advantage in studying these systems; for one, the possibility of direct experimental intervention in isolation to understand the intrinsic basis of rhythm. The other, is the tractable characterization of the rhythm in terms of burst-interval, cycle period, phase relationships that, by definition, repeat themselves.

We, too, have capitalized on this simple-most network capable of rich dynamical behavior. The central essence of rhythm generation, to be functionally relevant, is the regularity of the rhythm. We investigate how intrinsic and extrinsic noise affect the regularity of the switching of activity in two mutually inhibiting neurons. Switching in our model takes place due to the calcium-mediated potassium current (sAHP) build-up. The sAHP current allows one neuron to ‘escapes from inhibition’ of the other (Wang and Rinzel, 1992). Previous studies have investigated the role of noise in modulating neuronal firing rates (McDonnell and Abbott, 2009; Schmid et al., 2001; Wang, 1998; Nesse et al., 2008b,a). Current noise either reduces or enhances the gain of the firing frequency–current relationship depending on the type of intrinsic currents associated with the cell (Higgs et al., 2006). Noise is also seen to induce higher switching frequencies in CPGs of the rat spinal cord (Taccola, 2011). Some of the other consequences of extrinsic factors on the switching frequency of two mutually inhibiting neurons are described by Skinner et al.(Skinner et al., 1994). Our results demonstrate the non-monotonic effect of increasing external current on switching frequency. The initial decrease followed by the increase in switching frequency occurs because of the competing impact of increasing the external current (Figure 3); faster build-up of on sAHP current and higher-threshold acquired to escape from inhibition. Neuromodulators can modulate biophysical properties such as the channel conductance and calcium-binding rates (Schwartz et al., 2005) and can further modulate the range of switching repertoire of the network. It would be interesting to study, for example, the role of 5-HT in modulating the rhythm of the network (Kozlov et al., 2001).

We describe the effect of varying amplitude of current noise amplitude (Figure 4) on the switching dynamics. One of the most interesting insights from our study is that the switching frequency repertoire of the network is extended when current noise is introduced in the simulations (Figure 4). An analytical description of this phenomenon that uses a phenomenological neuron (Fitzhugh -Nagumo neuron) as an oscillatory system with multiple-separable timescales appears in Nesse et al. (Nesse et al., 2008a). We find that this noise-induced enhancement of the dynamical range of switching takes place at a small cost of an increase in variability. The CV of switching is also seen to depend on the amplitude of the external current stimulus (Figure 4C, A and D. Reliable switching in the presence of noise over a range of driving current indicates a match between the calcium spike frequencies driven by the current stimulus and the timescale sAHP over which sAHP integrates the current fluctuations. A similar mechanism via a match in timescales of fluctuations and sAHP current could explain the regular switching seen in many systems such as the Lamprey locomotion CPG and pre-Botzinger complex (Nesse et al., 2008b; Cangiano and Grillner, 2004). It may not always be possible to update a biological system’s intrinsic parameters in an activity-dependent manner. Here we show that external noise can be advantageous rather than a hindrance and extend the neuron’s dynamic range.

We examined the effect of calcium channel fluctuation, the most significant intrinsic noise source in neurons (Modchang et al., 2010). Forcing random failures in calcium spikes leads to the linear dependence of switching frequency on the failure rate (Figure 5). We also show that the rate of switching is invariant over multiple trials, i.e., switching frequency is insensitive to the position of failure of the calcium spike. This invariability over numerous trials demonstrates that sAHP current is a spike counter and can serve as a temporal integrator. Temporal integration has been implicated in audio and visual systems and involves collating spike patterns over time to improve detection or discrimination (Saija et al., 2019). A good temporal integrator requires that it maintain an average rhythm that is unaffected by input noise. The network with sAHP current can serve that purpose. A slow afterhyperpolarization that rises from *Na*^+1^*/K*^+1^ pump dynamics can also act as an integrator of spike number and serve as cellular memory on the time scales of the cycle periods of the locomotion rhythms (Pulver and Griffith, 2010). Separately, potassium current with slow inactivation has been implicated in modulating the synaptic plasticity and short term memory by changing the excitability of the cell (Turrigiano et al., 1996; Marder et al., 1996; Stackman et al., 2002). It would be interesting to investigate if sAHP dynamics simulated in our network would give rise to some form of cellular, short term memory.

The opening of calcium channels is rapid and closely follows the action-potential activity. However, as mentioned before, the response of the calcium-mediated potassium current, sAHP, is much slower (by order of magnitude). Thus each action potential leads to a fractional increase in the conductance of these channels. The firing ceases as the potassium current builds up over a train of action potentials. To account for fast calcium channel stochasticity and the slow cumulative increase sAHP, we modified the classical Gillespie algorithm (Gillespie, 1976b). We believe that the modified algorithm, TPG captures the fluctuations and progressions governing the neuronal and network dynamics over multiple timescales more accurately.

We also investigated the effect of varying calcium channel number while keeping the maximum conductance the same on the network’s switching dynamics (Figure 6). As expected, the CV of switching decreases as the channel number increases. However, the switching frequency has a non-monotonic dependence on the number of calcium channels. An upstroke in switching frequency is seen for a range of ion channel number (∼10 - 50). The larger fluctuations in the fraction of open channels result in this behavior. As the open channel numbers fluctuate widely, the network has more significant excursions through switching intervals dictated by the open fraction. The waiting times for transitions between states of calcium channels are too long, and higher frequencies of switching are not achieved when a small number of calcium channels are present (Figure 6 D). Interestingly, the number of channels that orchestrate the highest switching frequencies (∼50 - 300) are also realistic estimates for the number of L-type calcium channels present in the neuron. An increasing trend in switching frequency is seen again for large numbers of ion channels (*>* 500). It is noteworthy that the switching frequency at these high channel numbers (Figure 6 A for current = 14*µA/cm*^2^) is larger than the deterministic limit of switching frequency (Figure 3 D for current = 14*µA/cm*^2^). The small waiting times due to the large population statistics of calcium channels (Figure 6 D) lead to frequent fluctuations in the calcium current. A corollary insight from these calculations is that stochasticity also plays a role when channel numbers are large when slow dynamics are involved in contrast to the conventional belief. In sum, the competing effects of fluctuations and waiting times for calcium channels to change states lead to the non-monotonic behavior seen in switching frequency as we vary the number of calcium channels (Figure 6 A).

The brain is capable of generating regular firing patterns critical for several functions despite irregular inputs due to channel fluctuations, probabilistic neurotransmitter release diffusion of signaling molecules and probabilistic binding to receptors, etc. It is almost impossible to suppress all sources of noise experimentally. Computational modeling to isolate the consequences of noise to function is therefore valuable. Using a minimalistic model system for rhythm generation, our investigations have led to several novel insights into the contribution of noise to function. Each calcium fluctuation may not immediately affect the postsynaptic neuron; however, an ionic current like sAHP seems to keep an account of this miniature activity. We speculate that this may serve as a sub-cellular substrate of memory (Pulver and Griffith (2010)).

## AUTHOR CONTRIBUTIONS

Suhita Nadkarni (SN) conceived the project. SN and Subhadra Mokashe (SM) designed the model simulations. SM ran the simulations. SM and SN analyzed the data. SM and SN wrote the manuscript.

## ACKNOWLEDGMENTS

We would like to thank Dr. Collins Assisi for useful insights and past and present members of Nadkarni and Assisi labs for helpful discussions. SM would like to thank Dr. Satish Mokashe for helpful discussions and encouragement during the course of this project.

## CODE AVAILABILITY STATEMENT

The code is available at https://github.com/subhadram/insilico

